# Novel *EGLN1* variants identified in patients with erythrocytosis: a functional study

**DOI:** 10.64898/2026.06.12.731943

**Authors:** Aleša Kristan, Simon Fekonja, Nataša Debeljak

**Affiliations:** University of Ljubljana, Faculty of Medicine, Institute of Biochemistry and Molecular Genetics, Medical Centre for Molecular Biology, Ljubljana, Slovenia; Kemomed Ltd., Kemomed Research and Development, Ljubljana, Slovenia

**Keywords:** Erythrocytosis, Functional Analysis, *EGLN1* Gene, Variants of Uncertain Significance (VUS)

## Abstract

Erythrocytosis, a disorder with increased erythrocyte production, has a heterogeneous aetiology, including rare congenital types linked to dysregulation of the oxygen-sensing pathway. Variants in the *EGLN1* gene, encoding the prolyl hydroxylase that regulates hypoxia-inducible factor (HIF) stability, are associated with familial erythrocytosis type 3 (ECYT3). In patients with idiopathic erythrocytosis we previously identified two novel *EGLN1* variants, c.1072C>T (p.(Pro358Ser)) and c.1124A>G (p.(Glu375Gly)), classified as variants of uncertain significance. Herein, we performed in silico and in vitro analyses to assess their structural and functional effects, using the known pathogenic variant p.(His374Arg) as a positive control. AlphaFold3 predictions revealed minimal conformational changes in the protein core for all variants, while stability predictions suggested reduced protein stability. Functional assays in HEK293 cells demonstrated significantly decreased protein levels and stability for p.(Pro358Ser) and p.(Glu375Gly), comparable to p.(His374Arg). However, luciferase reporter assays showed that, unlike p.(His374Arg), the novel variants did not substantially impair EGLN1 enzymatic activity or activate HIF signalling. Our results suggest that the novel variants may contribute to erythrocytosis through destabilization of EGLN1, supporting further studies to elucidate their precise impact on hypoxia regulation. This study highlights the complexity of studying EGLN1 variants and the importance of functional evaluation for clinical interpretation.

## 1 INTRODUCTION

Erythrocytosis is a haematological disorder with an increased erythrocyte production, usually determined by an elevated haemoglobin and haematocrit in the blood picture.^1^ The major cause for increased mortality and morbidity in erythrocytosis patients are thrombotic events due to a higher blood viscosity.^2^ The aetiology for the disease is heterogeneous, with the secondary acquired causes and the primary acquired type Polycythaemia vera being the most common. On the other hand, the congenital erythrocytosis (also called familial erythrocytosis) is rare. In the majority of cases, despite the thorough investigation, the pathogenic cause for erythrocytosis is not resolved and patients remained idiopathic.^3, 4^

The precise regulation of oxygen homeostasis in the body is essential for normal body metabolism, with dysregulation potentially leading to several disease pathologies, including congenital erythrocytosis.^5, 6^ The major regulator of oxygen homeostasis are hypoxia-inducible factors (HIFs), that upregulates numerous targets genes involved in pathways that either promote oxygen delivery or decrease oxygen consumption, such as erythropoiesis, angiogenesis, iron metabolism, bone marrow microenvironment adjustments, and glucose metabolism.^5, 7, 8^ Transcription factor HIF is a heterodimer, consisting of a regulated alpha subunit (HIFα) and a constitutively expressed beta subunit (HIF1β, also called ARNT).^9^ The egl-9 family hypoxia-inducible factor 1 (EGLN1; also called prolyl hydroxylase domain-containing protein 2, PHD2) is from the family of dioxygenases that in the presence of oxygen, hydroxylates HIFα subunit. Hydroxylation mediates the binding of the von Hippel-Lindau disease tumour suppressor (VHL), leading to HIFα ubiquitination and subsequent protein degradation. Under low oxygen conditions (hypoxia), the EGLN1 activity decreases, which enables rapid accumulation of HIFα. Stable HIFα dimerises with beta subunit and bind to the hypoxia-response element (HRE) in the promoter of target genes, triggering their transcription.^10–12^ Three isoforms of HIFα are known, with HIF2α (official name endothelial PAS domain-containing protein 1, EPAS1) being the isoform strongly involved in erythropoiesis.^13, 14^ Also, three isoforms of EGLNs are known, among which EGLN1 seems to be critical oxygen sensor showing the highest oxygen-dependent activity.^15^

Variants in *EGLN1* are associated with familial erythrocytosis type 3 (ECYT3). ^16, 17^ To date, approximately 100 different germline *EGLN1* variants have been reported in patients with erythrocytosis, most of them were heterozygous. These include missense, nonsense, frameshift, duplication or deletion variants, which were dispersed throughout the *EGLN1*. In some patients, complications like thrombosis and hypertension occurred.^17–20^ In few cases, patients were also presented with rare neuroendocrine tumours, paragangliomas and phaeochromocytomas.^21, 22^ Functional studies employing diverse in vitro functional approaches demonstrated that some variants lead to either severe or subtle loss of EGLN1 function, reducing its ability to hydroxylate HIF.^19, 22–26^ Some studies also reported effects of variants on EGLN1 stability and splicing. ^19, 21^

In previous research we identified by targeted next-generation sequencing (NGS) two novel *EGLN1* variants, c.1072C>T (p.(Pro358Ser)) and c.1124A>G (p.(Glu375Gly)) in idiopathic erythrocytosis patients.^27^ Both variants had a predicted damaging effect by theoretical pathogenicity, were absent from the control populations of GnomAD, and were classified as variants of uncertain significance (VUS). Variant c.1072C>T (p.(Pro358Ser)) was also found in two affected family members, but the co-segregation evidence was not sufficient enough to upgrade its classification to likely pathogenic. The positioning of variants on protein structure revealed their location in the catalytic core, in the domain responsible for the hydroxylation reaction.^27^

In the present study, we performed further in silico and in vitro functional analyses to evaluate the effect of identified novel *EGLN1* variants c.1072C>T (p.(Pro358Ser)) and c.1124A>G (p.(Glu375Gly)) on the EGLN1 protein and on the subsequent development of the disease. A pathogenic variant c.1121A>G (p.(His374Arg)), with known effect on impaired EGLN1 function^22, 23^, was included in the study as a positive control.

## 2 MATERIALS and METHODS

### 2.1 In silico structural comparison analyses

The three-dimensional structures of EGLN1 wild-type (WT) and its variants p.(His374Arg), p.(Pro358Ser), and p.(Glu375Gly) were predicted using AlphaFold3, available at AlphaFold server (https://alphafoldserver.com/).^28^ Molecular visualization and structural analyses were performed with UCSF ChimeraX software.^29^ To assess conformational effects of variants, variant EGLN1 structures were superimposed onto the WT using the ChimeraX matchmaker tool. Sequence alignment was performed with the Needleman-Wunsch algorithm the BLOSUM-62 substitution matrix, applying the default iteration cut-off distance of 2 angstroms (Å) for structural fitting. Structural similarity was evaluated by pruned root-mean-square deviation (RMSD) values, which reflected differences only between well-aligned C*α* atoms retained during the iteration process, and overall RMSD values were also obtained.

### 2.2 In silico structural stability analysis

The effect of variants on protein stability was evaluated using web-based predictors DynaMut2 (https://biosig.lab.uq.edu.au/dynamut2/)^30^, mutation Cutoff Scanning Matrix (mCSM) (https://biosig.lab.uq.edu.au/mcsm/)^31^ and DUET (https://biosig.lab.uq.edu.au/duet/)^32^. All in silico predictions were performed using the WT EGLN1 structure generated by AlphaFold3. For each variant, predicted changes in Gibbs free energy (ΔΔG) were obtained, where negative values denote a destabilising effect and positive values indicate stabilisation of the protein structure.

### 2.3 Plasmids and cell culture for in vitro functional analysis

The novel *EGLN1* variants, p.(Pro358Ser)) and p.(Glu375Gly), together with previously reported pathogenic variant p.(His374Arg), were introduced into commercially available *EGLN1* expression plasmid (p.CMV6-EGLN1-MYC-DDK; OriGene Technologies) using the Q5 Site-Directed Mutagenesis Kit (New England Biolabs). Forward and reverse oligonucleotides used for in vitro mutagenesis are provided in **Supplementary table T1**. The presence of the inserted variants was validated by Sanger sequencing (Eurofins Genomics, Germany). For luciferase reporter assay commercially available *EPAS1* expression plasmid (p.CMV6-EPAS1-MYC-DDK; Origene Technologies), a Firefly luciferase reporter plasmid containing a hypoxia-response element (HRE) (pGL4.42; luc2P/HRE/Hygro; Promega), a Renilla luciferase control plasmid (pGL4.73; hRluc/SV40; Promega), and an empty vector (pCMV6-Entry, Origene Technologies) were also used.

Human embryonic kidney cells HEK293 were maintained in Minimal Essential Medium (MEM) (Sigma-Aldrich), supplemented with 10% foetal bovine serum (Sigma-Aldrich), 1% GlutaMAX (Gibco) and 1% antibiotic-antimycotic (Gibco). Cells were cultured under normoxic conditions (5% CO2, 37°C). Transient transfections were conducted using the PolyJet In Vitro DNA Transfection Reagent (SignaGen Laboratories).

### 2.4 In vitro analysis of EGLN1 protein levels

Protein levels of transfected *EGLN1* in HEK293 were analysed by seeding cells in 24-well plates, followed 24 h later by transfection with 500 ng of *EGLN1* plasmid carrying either WT or variant constructs. Cell lysates were prepared 24 h post-transfection using RIPA buffer. For immunoblots, 10 μg of protein lysates were separated by SDS-PAGE and transferred on membranes. Total proteins were labelled for normalization using the No-Stain Protein Labelling Reagent (Thermo Fisher). Immunodetection was performed with mouse anti-Myc primary antibody (clone 9E19, Origene) and goat anti-mouse secondary antibody (Jackson Immunoresearch). Immunoblots were imaged using the iBright FL 1500 (Thermo Fisher), and MYC signals were quantified and normalized to total proteins using iBright Analysis Software (Thermo Fisher).

### 2.5 In vitro analysis of EGLN1 protein stability

Protein stability was assessed using cycloheximide (CHX) chase assay. HEK293 cells were seeded in 12-well plates and transfected 24 h later with 500 ng of WT or variant *EGLN1* plasmids. At 24 h post-transfection, cells were treated with CHX (Sigma-Aldrich) at a final concentration of 100 µg/ml. Protein lysates were collected at 0 h, 3 h, 6 h, and 10 h after CHX addition and analysed by quantitative immunoblotting as described in section 2.4.

### 2.6 In vitro analysis of EGLN1 activity and HIF signalling

The functional effects of *EGLN1* variants on EGLN1 activity and further HIF signalling were assessed using an HRE-driven luciferase reporter assay. HEK293 cells were seeded in 24-well plates and, after 24 h, co-transfected with *EGLN1* WT (40 ng) or variant (40 to 115 ng) plasmid, *EPAS1* plasmid (200 ng), luciferase reporter plasmid (100 ng), luciferase control plasmid (10 ng), and empty vector to a total DNA input of either 350 ng or 540 ng. At 24 h post-transfection, cells were lysed with Passive Lysis Buffer (Promega) and luciferase activity was measured in 96-well plates using Dual-Glo Luciferase Reporter Assay (Promega Corporation) on a Synergy H4 microplate reader (Agilent BioTek). Protein lysates were analysed by immunoblotting as described in 2.4. Additional rabbit anti-EPAS1 primary antibody (Abcam), and goat anti-rabbit secondary antibody (Abcam) were used to detect EPAS1 signal.

### 2.7 Statistical analysis

All functional assays were performed in three independent experiments. Statistical analysis was conducted with GraphPad Prism Software (version 10.2.3. for Windows; GraphPad Software). For analysis of protein levels and luciferase reporter assays, statistical significance was determined by repeated measures one-way analysis of variance (ANOVA) under the assumption of sphericity, followed by Dunnett’s multiple comparison test. For CHX chase assay, statistical analysis was performed using a two-way repeated measures ANOVA with the assumption of sphericity, followed by Dunnett’s multiple comparison test. A p-value < 0.05 was considered statistically significant.

## 3 RESULTS

### 3.1 In silico comparison of wild-type and variant EGLN1 structures

The protein structures of EGLN1 WT and variants p.(His374Arg), p.(Pro358Ser), and p.(Glu375Gly) were predicted using AlphaFold3 and analysed in UCSF Chimera X. The core regions were predicted with high local confidence, but reliable predictions could not be obtained for the full-length WT and variant EGLN1 sequences, with a predicted template modelling score (pTM) of 0.62 or 0.63. Structural predictions for novel variants p.(Pro358Ser) and p.(Glu375Gly), as well as for known pathogenic variant p.(His374Arg), closely resembled WT in the core protein structure (**Figure 1**). Superimposition of WT and variant structures confirmed a high degree of similarity within well-aligned regions, with low pruned RMSD values (0.544 Å – 0.580 Å). While overall RMSD values were substantially higher (15.286 – 18.833 Å), likely due to low-confidence regions (**Supplementary figure S1**).

**Figure 1.**
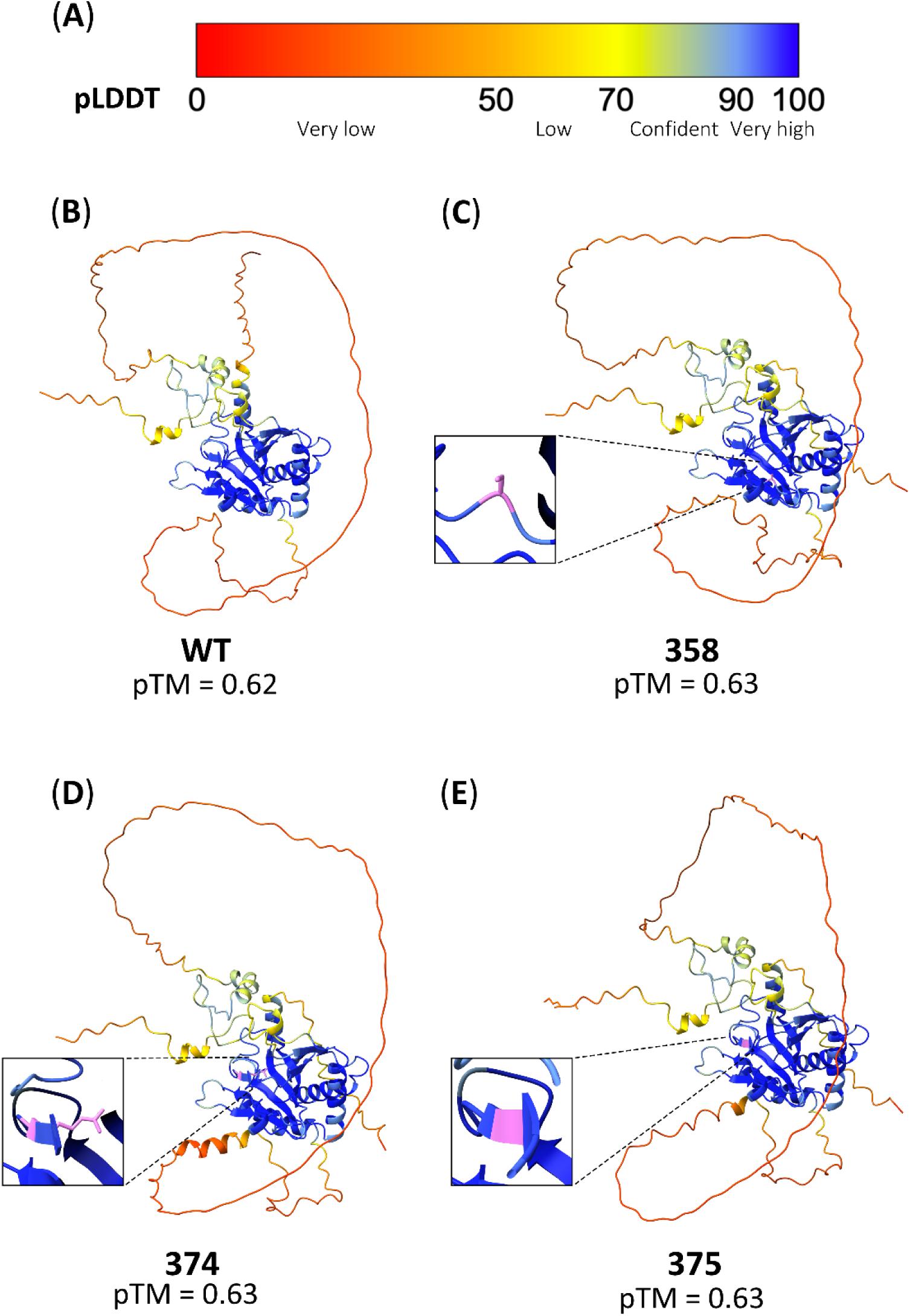
Predicted protein structures of EGLN WT and variants by AlphaFold3. (**A)** Colour scale of the predicted local distance difference (pLDDT) that indicates per-residue prediction confidence. (**B**) Structure of EGLN1 WT. (**C**) Structure of EGLN1 p.(Pro358Ser). (**D**) Structure of EGLN1 p.(His374Arg). (**E**) Structure of EGLN1 p.(Glu375Gly). Protein structures are visualized in UCSF Chimera X. The rectangle highlights the variant position, with purple indicating the variant amino acid. WT - wild-type; 358 - c.1072C>T (p.(Pro358Ser)); 374 - c.1121A>G (p.(His374Arg)); 375 - c.1124A>G (p.(Glu375Gly)); pTM predicted template modelling score.

### 3.2 In silico predictions of EGLN1 variants effect on structural stability

The effect on protein stability of the two novel EGLN1 variants, p.Pro358Ser and p.Glu375Gly, and the pathogenic variant p.His374Arg, was predicted with online servers DynaMut2, mCSM and DUET. As summarized in **Table 1**, all three computational tools consistently predicted a destabilizing effect for all variants. The DynaMut2 predictions are additionally illustrated in **Figure 2**, highlighting the structural impact of each mutation. These results suggest that the studied amino acid substitutions may compromise local stability and flexibility of EGLN1, potentially affecting its functional dynamics.

**Figure 2.**
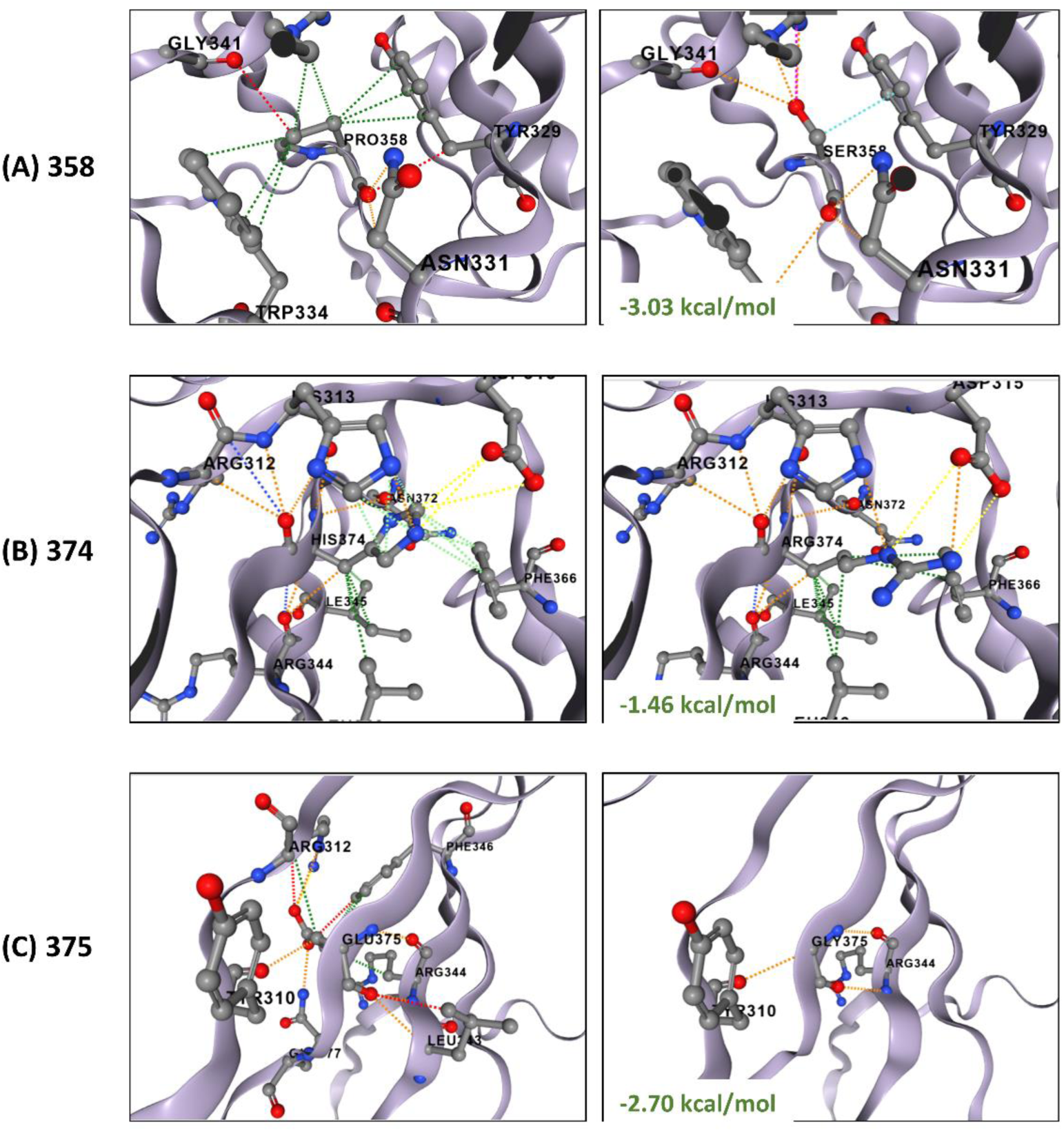
Predicted destabilising effects and structural impacts of EGLN1 variants using DynaMut2. (**A**) Stability and structural changes between EGLN1 WT and p.Pro358Ser variant. (**B**) Stability and structural changes between EGLN1 WT and p.His374Arg variant. (**C**) Stability and structural changes between EGLN1 WT and p.Glu375Gly variant. WT structures are presented in left panels and mutated structures in the right panels. WT and substituted residues, along with interacting amino acids, are displayed as sticks. Interactions, including aromatic (light green), hydrophobic (green), Van der Walls (turquoise), polar (orange), ionic (yellow) and hydrogen bonds (red) are represented by dashed lines. The predicted change in Gibbs free energy (ΔΔG) upon mutation, calculated by DynaMut2, is shown in green. 358 - c.1072C>T (p.(Pro358Ser)); 374 - c.1121A>G (p.(His374Arg)); 375 - c.1124A>G (p.(Glu375Gly)).

**Table 1.** Destabilising effects of EGLN1 variants on protein stability predicted by three in silico tools.

| Variant | DynaMut2 |  | mCSM |  | DUET |  |
| --- | --- | --- | --- | --- | --- | --- |
| | Predicted $\Delta\Delta G$ | Outcome | Predicted $\Delta\Delta G$ | Outcome | Predicted $\Delta\Delta G$ | Outcome |
| p.Pro358Ser | -3.03 kcal/mol | destabilising | -3.094 kcal/mol | highly destabilising | -2.926 kcal/mol | destabilising |
| p.His374Arg | -1.46 kcal/mol | destabilising | -2.249 kcal/mol | highly destabilising | -2.316 kcal/mol | destabilising |
| p.Glu375Gly | -2.70 kcal/mol | destabilising | -2.915 kcal/mol | highly destabilising | -2.975 kcal/mol | destabilising |

### 3.3 In vitro functional effect of EGLN1 variants on protein levels and stability

To assess the effect of EGLN1 variants on protein levels, MYC-tagged *EGLN1* WT and variant constructs were transfected into HEK293, and MYC signal was quantified by immunoblotting. As shown in **Figure 3A**, both novel EGLN1 variants, p.(Pro358Ser) and p.(Glu375Gly), had statistically significant lower protein levels than WT EGLN1 (p<0.0001). A similar decrease in protein levels was observed for the known pathogenic variant p.(His374Arg) (p<0.0001). Representative immunoblots are shown in **Supplementary figure S2.**

**Figure 3.**
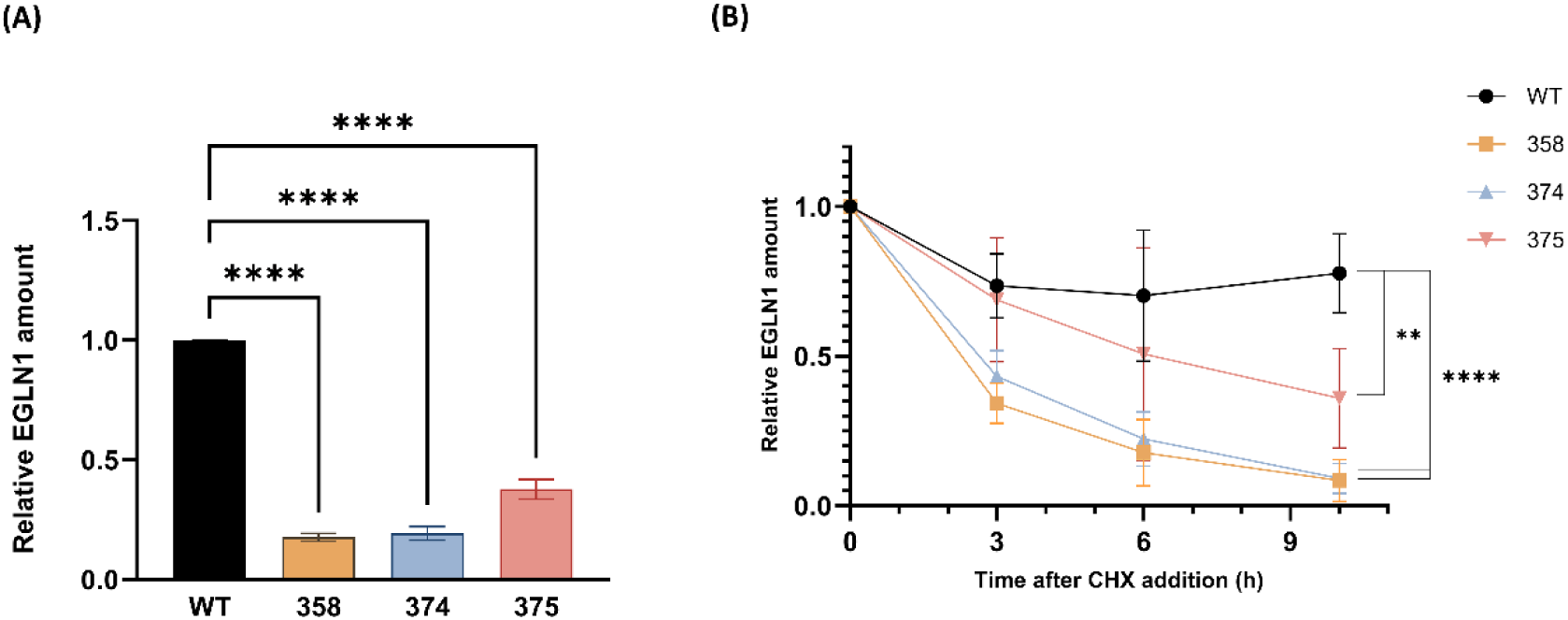
Decreased protein levels and stability of EGLN1 variants assessed by quantitative immunoblotting. **(A)** Study of EGLN1 protein levels by quantitative immunoblotting. For quantification of protein levels, MYC signal was analysed and normalized to total proteins. For representative immunoblot analysis see Supplementary figure S2. Data on the graph is presented as protein levels relative to the EGLN1 wild-type, and are mean ± standard error of three independent experiments. One-way ANOVA was used for the statistics. **(B)** Study of EGLN1 protein stability after addition of CHX by quantitative immunoblotting. For quantification, MYC signal was analysed and normalized to total proteins. For representative immunoblot analysis see Supplementary figure S3A. Graph shows protein levels measured at different time points after CHX addition relative to the initial levels at time 0 h. Data is presented as mean ± standard error of three independent experiments. Two-way ANOVA was used for the statistics. **** – p<0.0001; ** - p=0.0057; WT – wild-type; 358 - c.1072C>T (p.(Pro358Ser)); 374 - c.1121A>G (p.(His374Arg)); 375 - c.1124A>G (p.(Glu375Gly)).

To determine whether the decreased protein levels of EGLN1 variants p.Pro358Ser, p.Glu375Gly, and p.His374Arg were due to reduced protein stability, a CHX chase assay was performed, where CHX was added to stop protein synthesis. Protein levels were quantified on immunoblots at 0, 3, 6, and 10 h after CHX addition (**Supplementary figure S3A**). The results in **Figure 3B** show quantified protein levels at each time point, relative to their levels at time 0 h. WT EGLN1 protein remained relatively stable, with only a slight decline over time, consistent with the normal protein turnover. In contrast, all three variants p.Pro358Ser, p.Glu375Gly, and p.His374Arg, displayed progressively decreasing protein levels after CHX addition. At 10 h post-CHX treatment, all three EGLN1 variants showed a statistically significant decrease compared with WT (p<0.0001, p=0.0057 and p<0.0001, respectively) (**Figure 3B; Supplementary figure S3B**).

### 3.4 In vitro functional effect of variants on EGLN1 activity and HIF signalling

The effects of *EGLN1* variants on EGLN1 activity and downstream HIF pathway activation were evaluated by luciferase reporter assay (**Figure 4**). HEK293 cells were co-transfected with plasmids encoding *EGLN1* (WT or variants), *EPAS1* and a Firefly luciferase gene driven by HRE, followed by measurement of luciferase activity. Renilla luciferase was included to normalize for transfection efficiency (**Figure 4A**). Since EGLN1 variants showed reduced protein levels, *EGLN1* plasmid amounts were adjusted to achieve comparable protein levels across all constructs, thereby isolating the effect of variants on EGLN1 activity independent of protein abundance.

**Figure 4.**
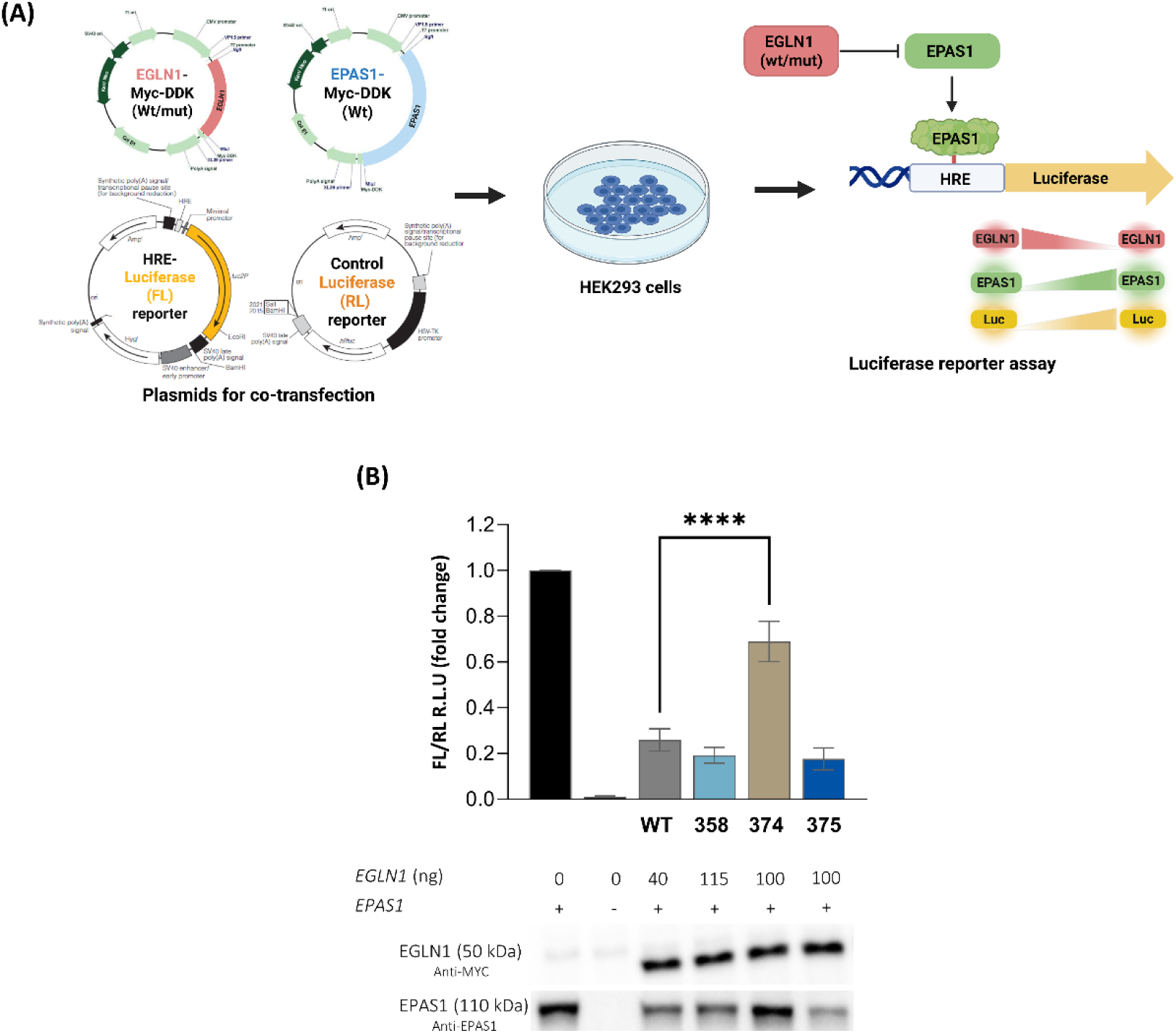
Lack of effect of novel *EGLN1* variants on EGLN1 activity and HIF signalling assessed by luciferase reporter assay. (**A**) Schematic overview of the luciferase reporter assay. (**B**) Luciferase activity measured after adjusting *EGLN1* plasmid amounts to achieve equal EGLN1 protein levels. Firefly luciferase signal was normalized to Renilla signal. Data are presented relative to the control without EGLN1 and shown as mean ± SD of three independent experiments. **** p<0.0001; WT – wild-type; 358 - c.1072C>T (p.(Pro358Ser)); 374 - c.1121A>G (p.(His374Arg)); 375 - c.1124A>G (p.(Glu375Gly)); HRE – hypoxia-response element; FL – Firefly luciferase; RL-Renilla luciferase; R.L.U – relative light unit; Luc – luciferase.

As shown in **Figure 4B**, addition of EGLN1 WT resulted in decreased EPAS1 protein levels and reduced luciferase activity compared with the control without EGLN1, consistent with EPAS1 degradation and consequent suppression of its transcriptional activity. Among the EGLN1 variants, only the known pathogenic variant p.(His374Arg) showed a statistically significant increase in luciferase activity compared with WT, consistent with its reported inhibition of EGLN1 activity (Ladroue et al., 2008, 2012). Immunoblots also confirmed increased EPAS1 protein levels for p.(His374Arg). Comparable results were obtained when equal plasmid amounts were transfected, indicating that novel variants p.(Pro358Ser) and p.(Glu375Gly) did not significantly alter luciferase activity compared with WT, despite reduced protein levels (**Supplementary Figure S4**).

## 4 DISCUSSION

High-throughput NGS technologies are now widely used to improve the diagnosis of rare inherited disorders, including hereditary erythrocytosis. ^33–35^ However, many identified variants remain classified as variants of uncertain significance (VUS), creating concern for carriers and dilemmas for clinicians in patient counselling.^35, 36^ The clinical relevance of VUS may be clarified through more extensive population data, family studies, and functional analysis.^36^ Here, we applied several in silico and in vitro functional assays to evaluate the effects of novel EGLN1 VUSs on protein conformation, abundance, stability and HIF signalling, and to explore their potential role in erythrocytosis.

In silico comparison of AlphaFold3 predicted structures indicated that both novel EGLN1 variants p.(Pro358Ser) and p.(Glu375Gly), as well as the pathogenic variant p.(His374Arg), have minimal effects on protein conformation in the highly confident core structure (**Figure 1**). This was further supported by pruned RMSD values below 1 Å (**Supplementary Figure S1**). Typically, RMSD values between 0–2 Å are observed for structures with a very high degree of sequence identity.^37^ However, reliance on pruned RMSD alone may underestimate global conformational changes. The substantially higher overall RMSD values observed, likely reflect deviations in regions of low-confidence predictions, rather than true conformational differences (**Supplementary Figure S1**). With more reliable full-length structures, the RMSD values might provide a more meaningful insight into structural similarity. Notably, the known pathogenic variant also showed a low pruned RMSD, suggesting that comparison of AlphaFold3-predicted structures alone currently cannot reliably capture the functional consequences of EGLN1 variant, and that functional assays remain essential for assessing their impact. Importantly, despite the low RMSD values suggesting minimal structural changes, in silico stability predictions consistently indicated that both novel, as well as the positive control variants destabilize protein stability (**Table 1**; **Figure 2**). Notably, the predicted destabilization for the novel variants was slightly more pronounced than that observed for the known pathogenic variant, as reflected by more negative values by all three predictors.

In vitro analyses first showed that both novel EGLN1 variants, p.(Pro358Ser) and p.(Glu375Gly), as well as the known pathogenic variant p.(His374Arg), had lower protein levels in HEK293 compared with WT EGLN1 (**Figure 3A**). Reduced protein levels for p.(His374Arg) have been reported previously.^22^ Based on in silico predictions, we assumed that decreased protein levels were due to a decrease in protein stability. This was confirmed by the CHX chase assay, which showed reduced stability for all three investigated EGLN1 variants p.Pro358Ser, p.Glu375Gly and p.His374Arg (**Figure 3B**). A similar loss of stability has been reported for other *EGLN1* variants identified in patients with idiopathic erythrocytosis and in patient with pheochromocytoma/paraganglioma and polycythemia.^19, 21^

The luciferase reporter assay demonstrated no substantial loss of EGLN1 activity or increased HIF signalling for the two novel EGLN1 variants, p.(Pro358Ser) and p.(Glu375Gly), as luciferase activity remained comparable to that of the WT (**Figure 4B**). In contrast, the known pathogenic variant p.(His374Arg) caused a robust increase in luciferase activity and EPAS1 protein levels, confirming previous reports of its strong impairment of EGLN1 activity.^22, 23^ Notably, when accounting for the reduced protein levels of the variants, the luciferase activity for p.(His374Arg) was comparable to that observed in the absence of EGLN1, whereas the novel *EGLN1* variants did not show a significant increase in luciferase activity (**Supplementary Figure S4**). Our findings suggest that novel variants p.(Pro358Ser) and p.(Glu375Gly) may cause only mild functional impairment, making their effects difficult to substantiate. The lack of substantial HIF activation despite reduced protein levels could also reflect sufficient residual EGLN1 activity or compensatory effects from endogenous EGLN1, potentially masking the impact of the transfected variants. Recent studies have highlighted the challenges of studying *EGLN1* variants, suggesting that some variants associated with congenital erythrocytosis appear to have subtle loss of function close to the WT protein.^19^ Despite its widespread use in EGLN1 functional studies^19, 22–25^, the luciferase reporter assay used here may have limited sensitivity for detecting subtle EGLN1 functional differences.

In summary, this study showed that the novel EGLN1 variants p.(Pro358Ser) and p.(Glu375Gly) reduce protein abundance and stability, similar to the known pathogenic variant p.(His374Arg). Unlike p.(His374Arg), the novel variants did not impair EGLN1 enzymatic activity or markedly activate HIF signalling under our luciferase reporter assay conditions, suggesting subtle functional effects or possible assay limitations. Further studies using more sensitive or complementary experimental approaches will help to clarify the contribution of variants p.(Pro358Ser) and p.(Glu375Gly) to hypoxia signalling and erythrocytosis.

## 5 AUTHOR CONTRIBUTIONS

A.K. and N.D. designed the study. A.K. wrote the manuscript. A.K. performed the laboratory experiments, in silico analysis and analysed the data. N.D., S.F. and A.K. reviewed and edited the manuscript. All authors approved the final version of the manuscript.

## ACKNOWLEDGEMENTS

We gratefully acknowledge Nuša Avguštinčič for assistance in performing the laboratory experiments. We acknowledge the use of ChatGPT for AI assisted text editing and grammar corrections. All content generated with its help was carefully reviewed and edited by the authors, who take full responsibility for the published content.

## 6 FUNDING INFORMATION

This study was supported by the Slovenian Research Agency, grant numbers L3-9279, L3-4511 and P1-0390, and the Young Researcher grant (to A.K.).

## 7 CONFLICTS OF INTEREST STATEMENT

S.F. is an employee of Kemomed Ltd. The authors declare no other conflicts of interest.

## 8 DATA AVAILABILITY STATEMENT

Datasets used and/or analysed during the current study are available from the corresponding author on reasonable request.

## 9 PATIENT CONSENT STATEMENT

Not applicable, as this study did not involve human participants.

## 10 CLINICAL TRIAL NUMBER

Not applicable, as this study was not conducted as a clinical trial.

## 11 ETHICS APPROVAL STATEMENTS

Not applicable, as this study did not involve human participants or animals.

## Notes

### Competing Interest Statement

The authors have declared no competing interest.

