## Supplemental data for "Novel *EGLN1* variants identified in patients with erythrocytosis: a functional study"

### Supplementary material

#### Supplementary table

##### Supplementary table T1: Oligonucleotides used for in vitro mutagenesis

| Variant | Oligonucleotide | Sequence 5' - 3' | Length (nt) |
|---|---|---|---|
| **c.1072C>T; p.(Pro358Ser)** | EGLN1_Q5_358_F | TGACATTGAA**T**CCAAATTTGATAGACTG | 28 |
| | EGLN1_Q5_358_R | GCAAACTGGGCTTTGCCT | 18 |
| **c.1121A>G; p.(His374Arg)** | EGLN1_Q5-374_F | CGCAACCCTC**G**TGAAGTACAA | 21 |
| | EGLN1_Q5-374_R | ACGGTCAGACCAGAAAAAC | 19 |
| **c.1124A>G; p.(Glu375Gly)** | EGLN1_Q5-375_F | AACCCTCATG**G**AGTACAACCAG | 22 |
| | EGLN1_Q5-375_R | GCGACGGTCAGACCA | 15 |

Mutated nucleotide in the oligonucleotide sequence is indicated with red. nt – number of nucleotides

### Supplementary figures

#### Supplementary Figure S1

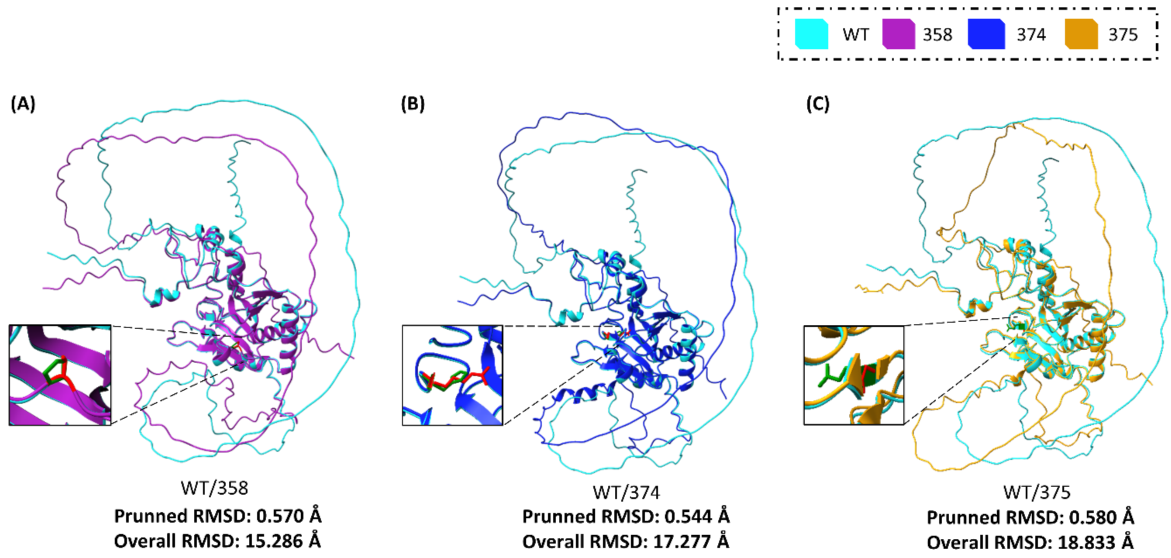


**Supplementary figure S1. High-similarity of EGLN1 WT and variant proteins in core regions assessed by ChimeraX. (A)** Superimposition of EGLN1 WT and p.(Pro358Ser). (**B**) Superimposition of EGLN1 WT and p.(His374Arg). (**C**) Superimposition of EGLN1 WT and p.(Glu375Gly). The rectangle highlights the variant position, with green indicating the WT amino acid and red indicating the mutant amino acid. RMSD values were obtained by ChimeraX software after superimposition. RMSD – root mean square deviation; Å- angstrom; WT - wild-type; 374 - c.1121A>G (p.(His374Arg)); 358 - c.1072C>T (p.(Pro358Ser)); 375 - c.1124A>G (p.(Glu375Gly)).

**Supplementary figure S2**

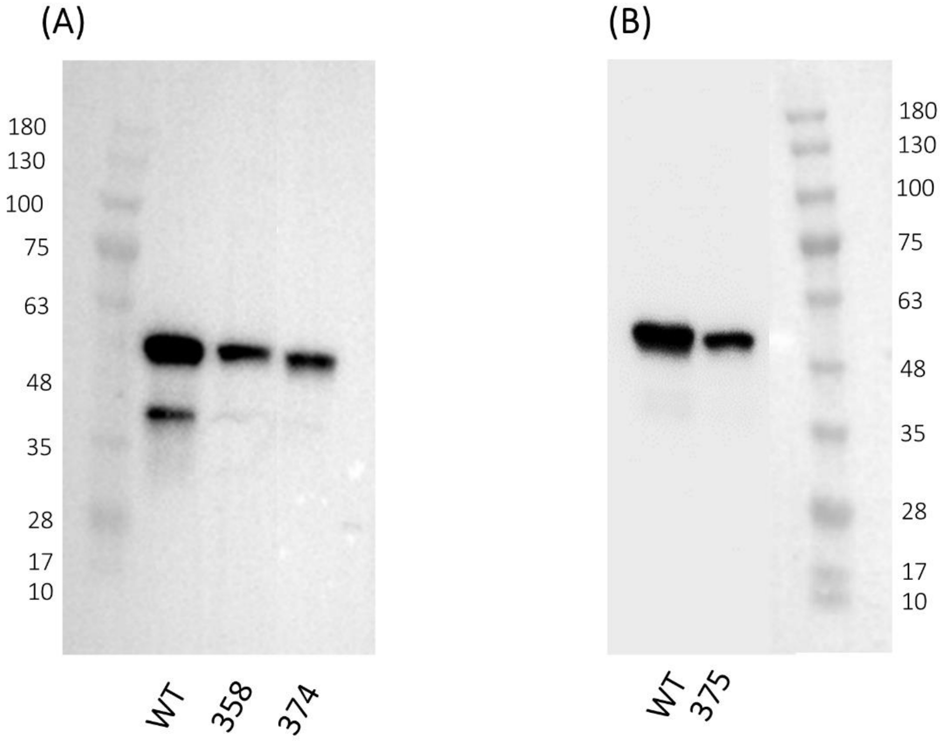


**Supplementary figure S2. Immunoblots showing EGLN1 WT and variants protein levels**
**A** Representative immunoblot for the EGLN1 WT, EGLN1 p.(Pro358Ser) and p.(His374Arg). **B** Representative immunoblot for the EGLN1 WT and EGLN1 p.(Glu375Gly). The signal shown on the immunoblots was obtained with anti-MYC primary antibody (Origene). For quantification of the amount of transfected EGLN1-MYC protein, anti-MYC signal was normalized to the labelled total proteins. Molecular weight marker was run along the samples; the numbers indicate kDa.

**Supplementary figure S3**

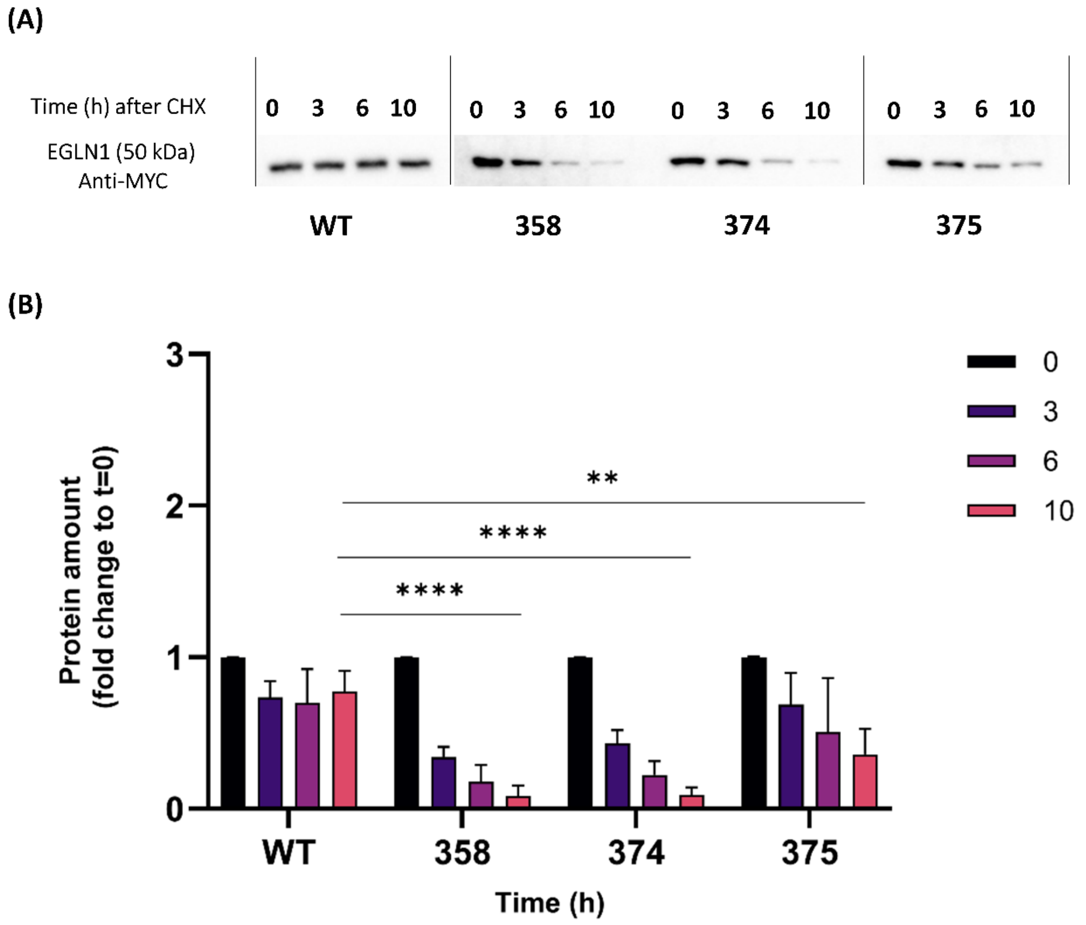


**Supplementary figure S3. Decreased stability of EGLN1 variants assessed by cycloheximide chase assay. A** Representative immunoblots of EGLN1 WT and variants p.(Pro358Ser), p.(His374Arg) and p.(Glu375Gly). Transfected EGLN1-MYC proteins were detected using anti-MYC antibody (Origene). **B** Quantification of protein amounts of EGLN1 WT and variants p.(Pro358Ser), p.(His374Arg) and p.(Glu375Gly) after treatment with cycloheximide (CHX). Quantified protein amounts on the graph are shown as relative to the initial levels at time 0 hours. For quantification of protein amount, MYC signal was analysed and normalized to labelled total proteins. Data is presented as mean ± standard error of three independent experiments. Two-way ANOVA was used for the statistics. ** - p=0.0057; **** – p<0.0001; WT – wild-type; 358 - c.1072C>T (p.(Pro358Ser)); 374 - c.1121A>G (p.(His374Arg)); 375 - c.1124A>G (p.(Glu375Gly)); CHX – cycloheximide.

**Supplementary figure S4**

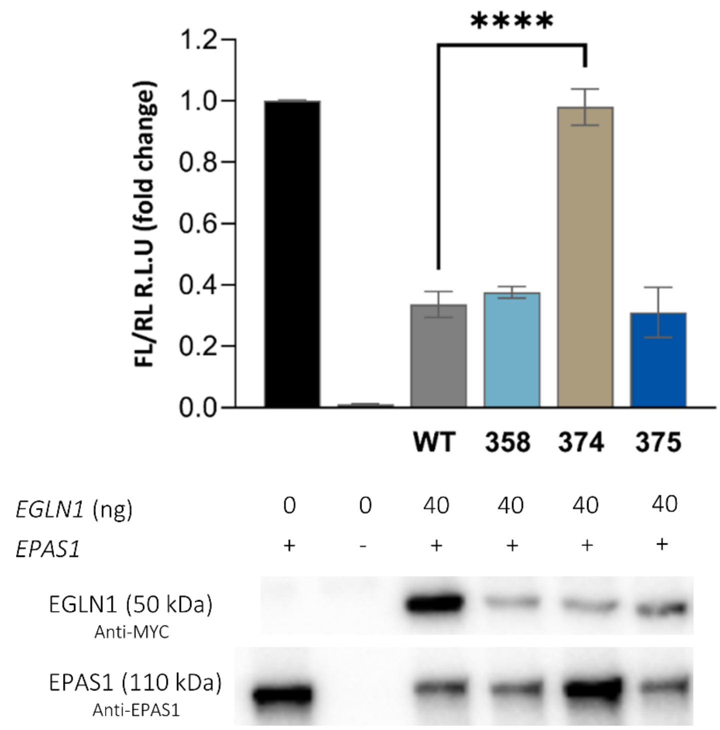


**Supplementary figure S4. Lack of effect of novel EGLN1 variants on EGLN1 activity and HIF signalling assessed by luciferase reporter assay with equal plasmid amounts.** Firefly luciferase signal was normalized to Renilla signal. Data are presented relative to the control without EGLN1 and shown as mean ± SD of three independent experiments. **** p<0.0001; WT – wild-type; 358 - c.1072C>T (p.(Pro358Ser)); 374 - c.1121A>G (p.(His374Arg)); 375 - c.1124A>G (p.(Glu375Gly)); FL – Firefly luciferase; RL- Renilla luciferase; R.L.U – relative light unit.
